# Inhibition of striatal dopamine release by the L-type calcium channel inhibitor isradipine co-varies with risk factors for Parkinson’s

**DOI:** 10.1101/2020.07.03.186411

**Authors:** Katherine R. Brimblecombe, Natalie Connor-Robson, Carole J. R. Bataille, Bradley M. Roberts, Caitlin Gracie, Bethan O’Connor, Rebecca te Water Naude, Gayathri Karthik, Angela J. Russell, Richard Wade-Martins, Stephanie J. Cragg

**Affiliations:** Department of Physiology, Anatomy and Genetics, University of Oxford, OX1 3PT, UK; Oxford Parkinson’s Disease Centre, University of Oxford, United Kingdom; Department of Chemistry, Chemistry Research Laboratory, University of Oxford, OX1 3TA; Department of Pharmacology, University of Oxford, OX1 3QT; Aligning Science Across Parkinson’s (ASAP) Collaborative Research Network, Chevy Chase, MD 20815, USA

**Author notes:** **Correspondence should be addressed to:** Dr Katherine Brimblecombe, Department of Physiology, Anatomy & Genetics, University of Oxford, OX1 3PT Oxford, UK. **Ethics statement** All procedures were licensed to be carried out at the University of Oxford under the UK Animals (Scientific Procedures) Act 1986. **Declarations of interest** The authors declare no competing interests. **Data availability** Data for all figures and code is available at: 10.5281/zenodo.7801868. Step-by-step experimental protocols are available at 10.17504/protocols.io.4r3l27dxxg1y/v1.

**Keywords:** Dopamine release, L-type calcium channel, dopamine transporter, Calb1, dopamine D2 receptor, striatum, Parkinson’s disease, sex differences, synuclein

## Abstract

Ca^2+^ entry into nigrostriatal dopamine (DA) neurons and axons via L-type voltage-gated Ca^2+^ channels (LTCCs) contributes respectively to pacemaker activity and DA release, and has long been thought to contribute to vulnerability to degeneration in Parkinson’s disease. LTCC function is greater in DA axons and neurons from substantia nigra pars compacta than from ventral tegmental area, but this is not explained by channel expression level. We tested the hypothesis that LTCC-control of DA release is governed rather by local mechanisms, focussing on candidate biological factors known to operate differently between types of DA neurons and/or be associated with their differing vulnerability to parkinsonism, including biological sex, α-synuclein, DA transporters (DATs), and calbindin-D28k (Calb1). We detected evoked DA release *ex vivo* in mouse striatal slices using fast-scan cyclic voltammetry, and assessed LTCC support of DA release by detecting the inhibition of DA release by the LTCC inhibitors isradipine or CP8. Using genetic knockouts or pharmacological manipulations we identified that striatal LTCC support of DA release depended on multiple intersecting factors, in a regionally and sexually divergent manner. LTCC function was promoted by factors associated with Parkinsonian risk, including male sex, α-synuclein, DAT, and a dorsolateral co-ordinate, but limited by factors associated with protection i.e. female sex, glucocerebrosidase activity, Calb1, and ventromedial co-ordinate. Together, these data show that LTCC function in DA axons, and isradipine effect, are locally governed and suggest they vary in a manner that in turn might impact on, or reflect, the cellular stress that leads to parkinsonian degeneration.

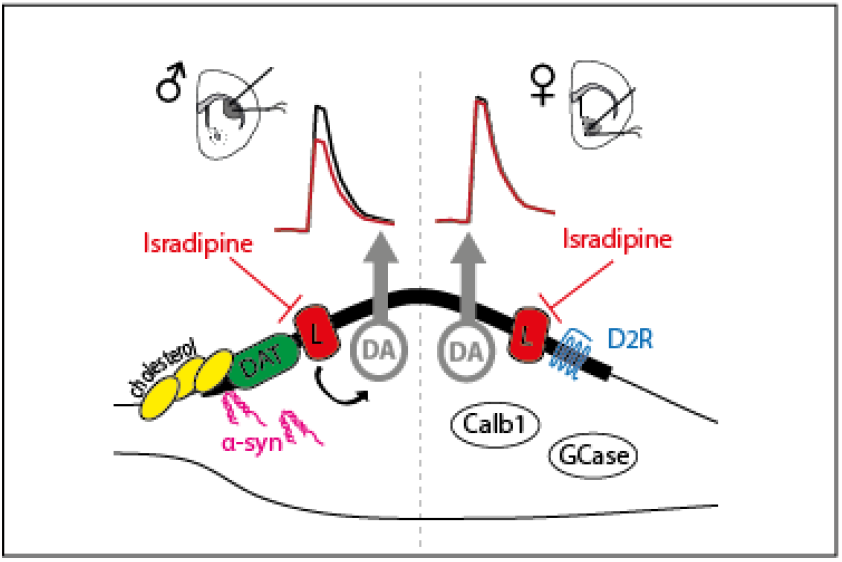

## Introduction

Ca^2+^ entry via voltage-gated Ca^2+^ channels (VGCCs) is important for pacemaker activity in dopamine (DA) neurons within the substantia nigra pars compacta (SNc) and critical for release of DA in the striatum (Guzman et al., 2009; Brimblecombe et al., 2015). However, Ca^2+^ also generates metabolic stress that promotes vulnerability of SNc DA neurons to degeneration in Parkinson’s disease (PD) (Surmeier et al., 2010). SNc DA neurons and their axons in dorsolateral striatum (DLS) are particularly vulnerable to parkinsonian degeneration, whereas DA neurons in ventral tegmental area (VTA) and their axons in ventral striatum (nucleus accumbens, NAc) are relatively spared. L-type VGCCs (LTCCs) support pacemaker activity, by facilitating ATP production in SNc (Zampese et al., 2022) and DA release in DLS, whereas in VTA and NAc, LTCCs do not operate strongly (Guzman et al., 2009; Brimblecombe et al., 2015; Zampese et al., 2022). However, this regional difference is not reflected by LTCC expression at the level of midbrain DA neurons where LTCC expression is greater in VTA (Poulin et al., 2014), suggesting that regulatory mechanisms locally govern function. Here, we explored factors that locally regulate the LTCC support of DA release from striatal DA axons in DLS vs NAc core.

Several biological factors and mechanisms that have direct relevance to the biology of DA neurons and/or to PD are candidates for LTCC regulators. On a population level, after aging, the biggest risk factor for PD is biological sex: men are twice as likely as women to develop PD (Wooten et al., 2004) but the biology underlying this difference is poorly understood. It has been established in cardiac myocytes that LTCC function is greater in males than females (Curl et al., 2008), but sex differences in LTCC function in central DA systems have not previously been explored. Here, we explored how LTCCs support DA release in both sexes of mice, and define sex differences.

Several genetic mutations and polymorphisms confer an increased risk to PD (Bekris et al., 2010). These range from rare, high penetrance mutations e.g. in the genes for α-synuclein (*SNCA*) and PINK-1/Parkin, to more common, less penetrant single-nucleotide polymorphisms, discovered by GWAS including *GBA* (gene encoding glucocerebrosidase, GCase) as the most commonly associated genetic risk factor. Interestingly, variants in *SNCA* occur as both rare, high penetrance and common, low penetrance forms. Here, we tested whether α-synuclein or GCase levels could impact on LTCC support of DA release.

We also tested whether several known physiological regulators of DA output from SNc and VTA DA neurons might support a differential contribution of LTCCs to DA release. We focussed on calbindin-D28K (Calb1), the DAT, and DA D_2_-receptors (D_2_Rs), each of which have been reported to interact with LTCCs in different systems. Calb1 is enriched in PD-resistant VTA neurons (Gerfen et al., 1987; Poulin et al., 2014), limits DA release in NAc (Brimblecombe et al., 2019) and, in cellular expression systems, can bind to and limit function of LTCCs (Lee et al., 2006). We report here that Calb1 limits LTCC support of DA release in a regionally and sexually divergent manner. DATs are enriched in SNc and VTA DA neurons, with highest expression levels reported in SNc, in both mouse and human brains (Ciliax et al., 1995; Poulin et al., 2014), and differential DAT expression has long been hypothesised to be a driving factor in sensitivity to PD on cellular and population levels (Ritz et al., 2009; Lohr et al., 2017). In cellular expression systems, DATs and other electrogenic monoamine transporters promote LTCC currents (Cameron et al., 2015). We report here that DATs play a role in facilitating how LTCCs support axonal DA release. Lastly, D2Rs are described as more enriched in SNc than VTA DA neurons (Greene et al., 2005; Poulin et al., 2014), and can inhibit LTCC function (via βγ-inhibition) in striatal GABAergic projection neurons (Hernández-López et al., 2000; Olson et al., 2005). Despite D2Rs not being associated with either risk or protection in PD (not least due to their broad expression pattern), they are well placed to modulate LTCC support of DA release. We report here that D2Rs operate a limiting factor on LTCC function in NAc. Overall, we reveal a network of factors that, by either inhibiting or facilitating LTCC function in DA axons, contribute to a greater role for LTCCs in supporting DA release in DLS than NAc, and in males than females, and a correspondingly varying outcome of LTCC inhibitor isradipine on DA release.

## Materials and methods

### Animals

We used male and female adult mice aged 3-10 months. Wild-type mice were C57Bl6/J (Charles River). Calbindin-knockdown (CalbKD) animals were generated by crossing homozygous DAT^IRES*cre*^ (https://www.jax.org/strain/006660) mice with homozygous Calb_TM2_ mice (loxP sites flanking Exon1) (Barski et al., 2002) donated by Prof. Meyer (University of Munich) (available from Jax https://www.jax.org/strain/031936 and EMMA: strain name B6.(R1)-Calb1tm2Mpin), generating double-heterozygous offspring with decreased calb1 expression in DAT-expressing cells as previously published (Brimblecombe et al., 2019). CalbWT control mice were age- and sex-matched heterozygous DAT^IRES*cre*^ mice. *SNCA*-OVX and *Snca*-null mice were generated in house as previously published (Janezic et al., 2013) (OVX https://www.jax.org/strain/023837; null https://www.jax.org/strain/003692). *Gba*^+/-^ mice were purchased from Jax (https://www.jax.org/strain/002594).

### Slice preparation

Mice were killed by cervical dislocation, the brains removed, and 300 μm coronal striatal slices prepared as described previously (Brimblecombe et al., 2015) in ice-cold HEPES-based buffer saturated with 95% O_2_/5% CO_2_, containing in mM: 120 NaCl, 20 NaHCO_3_, 6.7 HEPES acid, 5 KCl, 3.3 HEPES salt, 2 CaCl_2_, 2 MgSO_4_, 1.2 KH_2_PO_4_, 10 glucose. Slices were incubated at room temperature for ≥ 1 hour in HEPES-based buffer before experiments. All procedures were licensed to be carried out at the University of Oxford under the UK Animals (Scientific Procedures) Act 1986.

### Fast-scan cyclic voltammetry (FCV)

*Ex vivo* DA release was monitored in acute coronal slices using fast-scan cyclic voltammetry (FCV) as previously described (Brimblecombe et al., 2015). Slices were superfused in a recording chamber with bicarbonate-buffered artificial cerebrospinal fluid (aCSF) saturated with 95%O_2_/5%CO_2_ at 31–33 °C, containing in mM: 124 NaCl, 26 NaHCO_3_, 3.8 KCl, 0.8-3.6 CaCl_2_ (as stated), 1.3 MgSO_4_, 1.3 KH_2_PO_4_, 10 glucose. Evoked extracellular DA concentration ([DA]_o_) was monitored by FCV using a Millar voltammeter (Julian Millar, Barts and the London School of Medicine and Dentistry) and single-use carbon-fibre microelectrodes (7-10 μm diameter) fabricated in-house (tip length 50-100 μm). A triangular voltage waveform (range -700 mV to +1300 mV vs. Ag/AgCl) was applied at 800 V/s at a scan frequency of 8 Hz. Electrodes were switched out of circuit between scans. Electrodes were calibrated using 2 μM DA, prepared immediately before calibration using stock solution (2.5 mM in 0.1M HClO_4_ stored at 4 °C). Signals were attributed to DA due to the potentials of their characteristic oxidation (500-600 mV) and reduction (-200 mV) peaks. Step-by-step methods can be found at protocols.io: 10.17504/protocols.io.4r3l27dxxg1y/v1

### Electrical stimulation

DA recordings were obtained from dorsolateral quadrant of dorsal striatum, the dorsolateral striatum (DLS), or nucleus accumbens core (NAc). DA release was evoked by a local bipolar concentric Pt/Ir electrode (25 μm diameter; FHC inc. ME, USA) placed approximately 100 μm from the recording electrode. Stimulus pulses (200 μs duration) were given at 0.6 mA (perimaximal in drug-free control conditions). Stimulations were single pulses (1p) were repeated at 2.5 minute intervals. A stable pre-drug baseline of 6 stimulations before isradipine or CP8 was washed on for 12 stimulations. A stable baseline was determined during the experiment when the average peak oxidation for stimulus 5,6 was 95-105% of stimulus 1 and 2. All data were obtained in the presence of the nAChR antagonist, dihydro-β-erythroidine (DHβE, 1 μM) to exclude the strong effects of nAChRs on DA release (Zhou et al., 2001; Rice and Cragg, 2004; Cachope et al., 2012; Threlfell et al., 2012).

### Drugs and solutions

Reagents were purchased from Torcis or Sigma Aldrich (cocaine, water-soluble cholesterol). Stock solutions were made to 1000-2000x final concentrations in H_2_O (DHβE, cocaine), DMSO (isradipine, L-741,626, CP8), 0.1 M HCl (nomifensine) or ethanol (lidocaine) and stored at -20°C. Drugs were diluted to their required concentrations in aCSF immediately prior to use. Drug concentrations were chosen in accordance with previous studies (Acevedo-Rodriguez et al., 2014; Brimblecombe et al., 2015). Water soluble cholesterol or vehicle control Me-ß-cyclodextrin were dissolved in 10 ml aCSF, and slices were incubated and oxygenated for 30 mins before transferring to recording chamber as previously described (Threlfell et al., 2021).

### CP8 synthesis

According to the method (Kang et al., 2013) 2-(3-chlorophenyl)ethylamine (357 μL, 2.58 mmol, 1.0 eq.) was added in one portion to a solution of cyclopentane isocyanate (290 mg, 2.58 mmol, 1.0 eq.) in CH2Cl2 (10 mL) and the reaction was stirred at room temperature for 5 h (completion was monitored by LRMS). Malonyl chloride (276 μL, 2.84 mmol, 1.1 eq.) was then added dropwise under vigorous stirring over 5 min. The resulting mixture was stirred for 1 h and concentrated *in vacuo*. The residue was purified by column chromatography on silica gel (EtOAc/pentane, 1:4) to afford the title product as a white solid (687 mg, 80%). 1H NMR (400 MHz, CDCl3) δ 7.25 – 7.19 (m, 3H), 7.13 (dt, *J* = 6.7, 2.1 Hz, 1H), 5.20 – 5.07 (m, 1H), 4.13 – 4.03 (m, 2H), 3.62 (s, 2H), 2.92 – 2.84 (m, 2H), 1.94 (tqd, *J* = 8.0, 4.9, 2.0 Hz, 4H), 1.89 – 1.79 (m, 2H), 1.61 – 1.57 (m, 2H); 13C NMR (101 MHz, CDCl3) δ 164.9, 164.6, 151.1, 140.0, 134.5, 123.0, 129.2, 127.3, 127.1, 54.6, 42.8, 40.3, 33.8, 28.8, 25.7; LRMS m/z (ESI^-^) 333.1 [M-H]^-^. The data were in accordance with those reported in the literature (Kang et al., 2013)

### Data and statistical analysis

FCV data were acquired and analysed using Axoscope 10.5 (Molecular devices) and Excel macros written locally. Data are expressed as mean ± standard error of the mean (SEM), and N = number of slices (typically 1 per animal) from at least 3 animals. Data from each slice were obtained by averaging 4 recordings prior to drug wash-on (control) vs. average of 4 recordings after 20 minutes. Typically data are shown normalised to control conditions to standardise for variation in initial evoked extracellular DA concentrations ([DA]_o_). Population means were compared using one-or two-way ANOVA with Dunnett’s post-test and unpaired t-test where appropriate using GraphPad Prism (9.3.1). Curve fits were performed in GraphPad Prism. Data for all figures are available 10.5281/zenodo.7801868.

### Classification tree

A classification tree was performed in R(4.2.1) (10.5281/zenodo.7801868). The normalised effect of isradipine for each individual experiment was annotated with the experimental condition: sex and genotype of mouse and presence of drug prior to washing on isradipine (fig5 data tab in xlsx file 10.5281/zenodo.7801868). These data were sorted using a classification tree using Gini impurity algorithm with a cut off of 0.91 normalised effect of isradipine, which corresponds to 2 s.d. above the mean effect of isradipine in DLS of male WT mouse (i.e. the original recording conditions where we identify a significant effect of isradipine on decreasing DA (Brimblecombe et al., 2015). It should be noted that 0.91 is also 2 s.d. below mean normalised effect of isradipine in male WT NAc (2.4 mM [Ca^2+^]_o_), which was our original recording conditions when we did not identify an effect of isradipine). Due to the structure of the classification tree whereby, factors were effectively sorted with no outliers (i.e. the leaves (final nodes) were mostly pure with probabilities of 0 or 1), we determined it was unsuitable to run a logistic regression analysis on these data.

### Western blots

Mice were culled by cervical dislocation, brains extracted and dissected on ice. The striatum and midbrain were extracted from both hemispheres and snap frozen. Samples were prepared for western blot as previously described (Connor-Robson et al., 2019). Briefly, tissue was homogenised on ice in RIPA buffer containing protease inhibitors (Roche) and protein concentration determined by BCA assay. Following dilution Laemmli buffer was added and samples were boiled for 5 minutes at 95°C. Samples were loaded and run on 4-15% criterion-TGX gels and transferred to PDVF membranes (BioRad). Membranes were blocked for 1 hour at room temperature and then probed with primary antibody (calb1: 1:1000 Cell Signalling #13176) overnight at 4°C. After incubation with HRP conjugated secondary antibodies the blot was developed using chemiluminescent HRP substrate.

## Results

### Sex difference in the LTCC support of DA release

We and others, have to date principally explored LTCC function in males rather than females (Guzman et al., 2009, 2018; Ilijic et al., 2011; Brimblecombe et al., 2015). There is a ∼2:1 prevalence of PD in males and females and it is therefore important to understand the biology of LTCCs in both sexes, in order to gain a fuller understanding of disease aetiology but also drug action. Here, we tested whether LTCCs support DA release in DLS of female mice, which they do in male mice (Brimblecombe et al., 2015). We first assessed DA release and uptake parameters. We found no difference in peak evoked extracellular concentrations of DA ([DA]_o_) evoked by single electrical stimulus pulses (1p) (**Figure 1A,B**, t-test t_14_=0.14 *P=0*.*89*), but did find faster DA clearance rates in females (**Figure 1A,C**, T-test with Welch’s correction t_29.9_=6.59 *P<0*.*0001*), consistent with greater DAT levels in the striatum of female mice (Costa et al., 2021). We then tested the effect of LTCC inhibitor isradipine (5 μM) on 1p-evoked [DA]_o_ in DLS and confirmed a decrease in evoked [DA]_o_ in male mice (**Figure 1D**, comparison of normalised peak [DA]_o,_ One-sample t-test t_2_=5.6, *P*=0.03, N=3) as published previously (Brimblecombe et al., 2015) but, by contrast, not in female mice (**Figure 1D**, comparison of normalised peak [DA]_o,_ One-sample t-test t_4_=0.87, *P*=0.44, N=5), revealing a statistically significant difference between the two sexes (**Figure 1E**, Two-way ANOVA: sex x drug interaction, F_16,67_=3.0, *P*<0.001; main effect of sex, F_1,67_=44.9, *P*<0.001).

**Figure 1.**
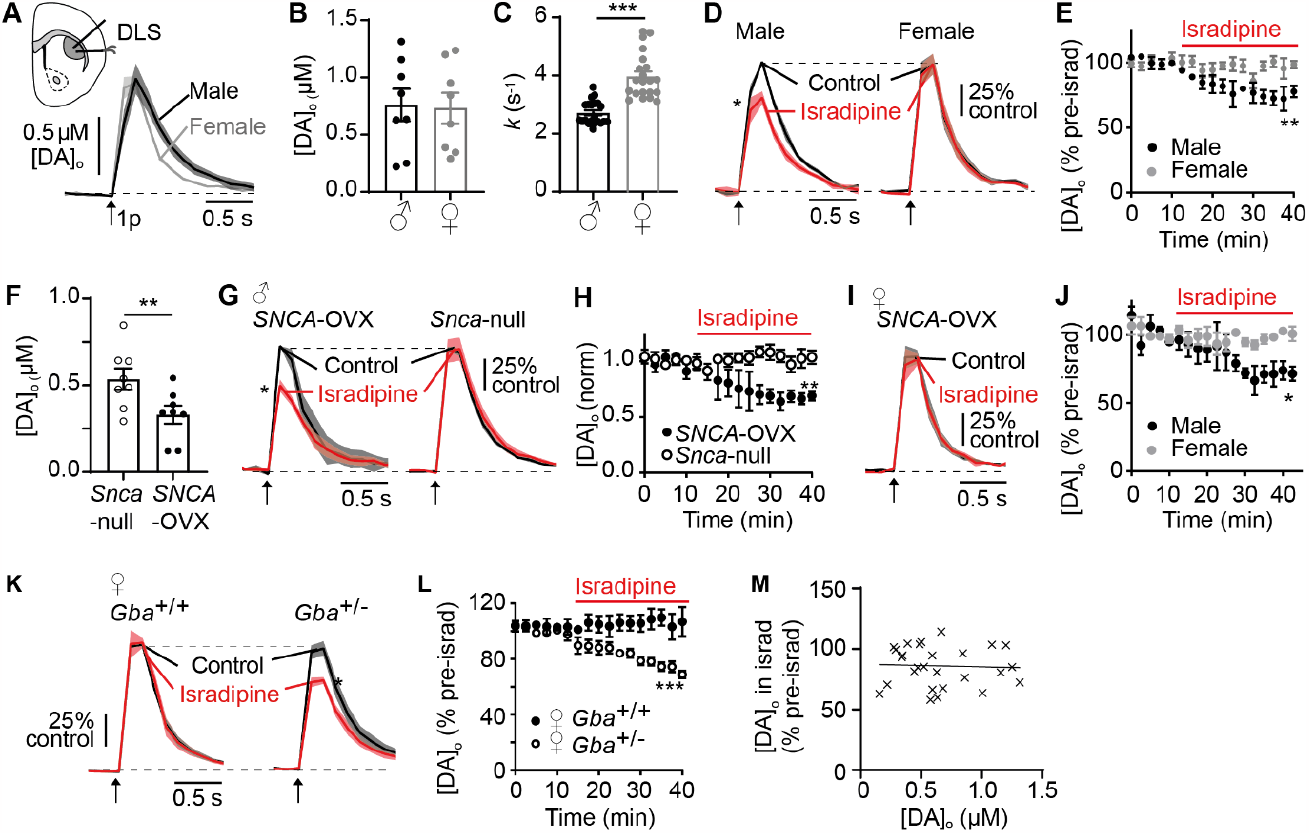
Male sex, presence of α-synuclein, and knockdown of GBA promote LTCC support of DA release in DLS. (**A**) Mean [DA]_o_ ± SEM vs time evoked by a single pulse (1p, *arrow*) in DLS of male (*black*) versus female (*grey*) wild-type mice. (**B**) Summary of peak evoked [DA]_o_ ± SEM, N = 6 mice. (**C**) Decay constants of falling phases of 1p-evoked DA transients in DLS of male versus female wild-type mice. Data from N = 12 transients from 6 mice. (**D**) Mean transients for [DA]_o_ ± SEM vs time evoked by 1p (*arrow*) in control conditions (*black*) versus isradipine (5 μM, *red*) in male (*left*) and female (*right*) normalised to own control. (**E**) Peak [DA]_o_ ± SEM as % of pre-isradipine control levels versus time. N = 3 male, 5 female. (**F**) Summary of peak [DA]_o_ evoked by 1p in DLS from male *SNCA*-OVX vs *Snca*-null mice. N = 8 mice. (**G**) Mean [DA]_o_ ± SEM vs time evoked by 1p (*arrow*) in control versus isradipine in *SNCA*-OVX (*left*) and *Snca*-null (*right*), normalised to control. (**H**) Peak [DA]_o_ ± SEM as % of pre-isradipine control levels vs time. N= 4 mice. (**I**) Mean [DA]_o_ ± SEM vs time evoked by 1p (*arrow*) in control and isradipine in female SNCA-OVX mice, normalised to control. N = 3 mice. (**J**) Peak [DA]_o_ ± SEM as % of pre-isradipine control levels vs time of female vs male *SNCA*-OVX mice. (**K**) Mean [DA]_o_ ± SEM vs time evoked by 1p (*arrow*) in DLS in control and isradipine in female wild-type (*Gba*^+/+^) versus GBA-het (*Gba*^+/-^) mice, normalised to control. (**L**) Peak [DA]_o_ ± SEM as % of pre-isradipine control levels vs time for female *Gba*^+/+^ (N=3) versus *Gba*^+/-^ mice (N=5). (**M**) Peak [DA]_o_ in isradipine (as % of pre-isradipine control levels) versus peak [DA]_o_ (μM) in pre-isradipine control conditions plotted for all data shown in conditions in A-L. *< 0.05 ** <0.01 ***<0.001.

### Genetic contributors to PD promote LTCC support of DA release

Mutations in the *SNCA* gene including common, lower penetrance SNPs identified by GWAS (Alegre-Abarrategui et al., 2019) are associated with rare monogenic forms of PD, while mutations in *GBA* are the most common genetic factor in PD. We tested whether α-synuclein facilitates LTCC support of DA release, using a BAC transgenic mouse line that overexpresses human wild-type α-synuclein (*SNCA*-OVX) (Janezic et al., 2013), versus the background control line that are null for mouse α-synuclein (*Snca*-null). Mean peak [DA]_o_ evoked by 1p in DLS of *SNCA*-OVX mice were ∼30% less than *Snca*-null mice (**Figure 1F**, unpaired t-testt_14_=2.55 *P*= 0.023) as seen previously (Janezic et al., 2013; Roberts et al., 2020; Threlfell et al., 2021). In DLS of male *SNCA*-OVX mice, isradipine decreased 1p-evoked [DA]_o_ by ∼30%, (**Figure 1G**, comparison of peak [DA]_o_ t_3_=6.6 *P*=0.007 N=4) as seen in WT (**Figure 1C**) (Brimblecombe et al., 2015)). However, in male *Snca*-null mice, isradipine had no effect on 1p-evoked [DA]_o_ (**Figure IG**, comparison of normalised peak [DA]_o_ t_3_=0.25 *P=*0.82 N=4), showing a statistically significant difference between the two genotypes (**Figure 1H**, two-way ANVOA: genotype X drug interaction, F_17,102_=3.0, *P*=0.0003; main effect of genotype, F_1,6_=4.89 *P*=0.002) and suggesting a dependence of LTCC function on α-synuclein.

We did not see an LTCC role in DA release in females in WT mice (see Figure 1C), but tested whether LTCC support of DA release in females could be established in either of two different PD-related mouse lines. Firstly, we explored whether overexpression of α-synuclein could result in LTCC support of DA release. However, in female *SNCA*-OVX mice, isradipine did not decrease [DA]_o_ in DLS (**Figure 1I**, comparison of normalised peak [DA]_o_ t_2_=2.2 *P*=0.16), indicating a significant difference between male and female *SNCA*-OVX mice (**Figure 1J**, two-way ANVOA: genotype X drug interaction, F_17,102_=1.89, *P*=0.027; main effect of genotype, F_1,6_=7.85 *P*=0.03). Secondly, since mutations in the GCase *GBA* gene are associated with PD and seen more commonly in women with PD than men (Li et al., 2021) we explored whether LTCC support of DA release varied with GCase levels in females, by comparing mice that were heterozygous for the mouse *Gba* gene *(Gba*^*+/*-^) with wild-type littermate homozygous controls (*Gba*^+/+^). Isradipine did not modify 1p-evoked [DA]_o_ in DLS *Gba*^+/+^ female controls, as seen for other wild-type females, but strikingly, isradipine did inhibit 1p-evoked [DA]_o_ in female *Gba*^+/-^ mice (**Figure 2K**, comparison of normalised peak [DA]_o_, *Gba*^+/+^(WT), t_2_=0.33 *P*=0.77; *Gba*^*+/*-^, t_4_=12.03 *P*=0.0003), showing a significant difference between these two genotypes (**Figure 1L:** two-way ANVOA: genotype X drug interaction F_16,96_=6.4 *P*<0.0001; main effect of genotype, F_1,6_=16.2 *P*=0.007). These observations indicate that a role for LTCCs in supporting DA release is exposed when GCase function is compromised.

**Figure 2.**
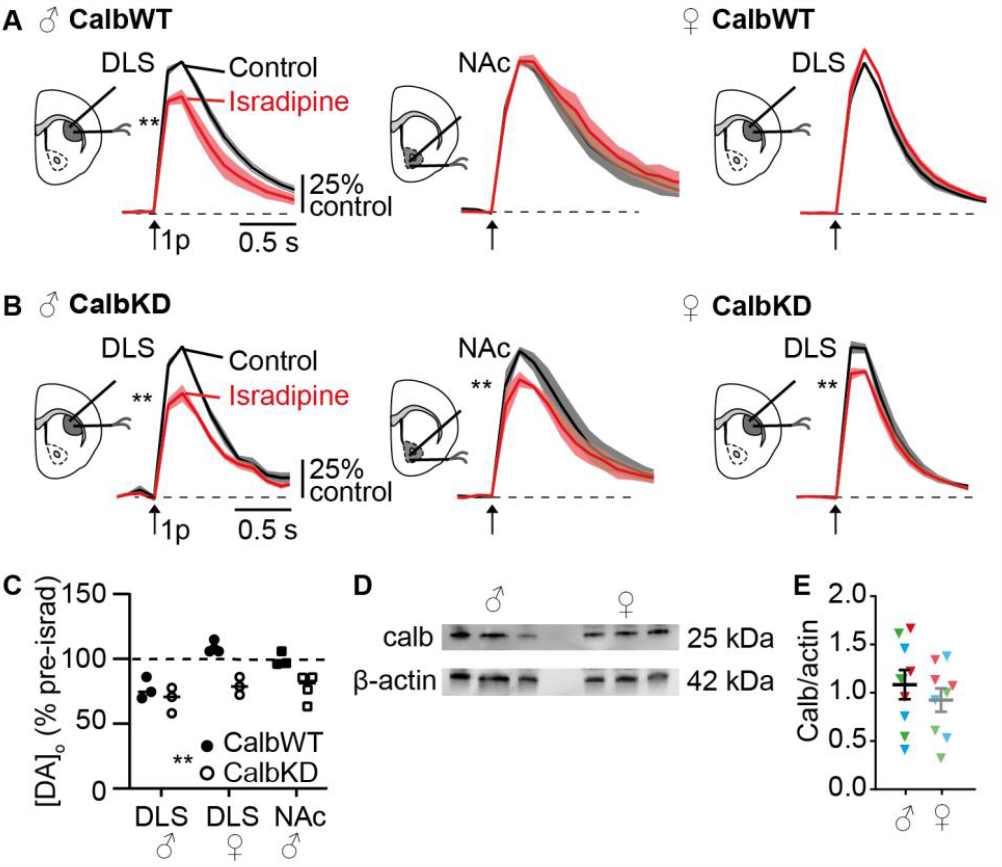
Calb1 limits LTCC function in NAc in male mice, and in DLS in female mice. (**A,B**) Mean [DA]_o_ ± SEM vs time evoked by single pulses (1p, *arrow*) in control and isradipine (5 μM, *red*) in (A) CalbWT mice or (B) CalbKD mice, in male DLS (*left*), male NAc (*centre*), and female DLS (*right*) normalised to their own control conditions. ** *P<*0.01 comparison of peak [DA]_o_(**C**) Summary of peak [DA]_o_ remaining after isradipine (% of pre-isradipine control) in CalbWT and CalbKD striatum. ** *P<*0.01 Effect of genotype 2-way ANOVA. N= 3 pairs DLS male and female N=4 NAc (**D**) Representative examples of Western blots for 3 male and 3 female showing actin (42 kDa) and Calb1 (25 kDa). (**E**) Calbindin-D28K/β-actin protein ratios in male and female striatum. Mean ± SEM; colours group technical replicates (i.e blue points correspond to blot shown in C, and are samples from which stats correspond to, N=3 brains, repeated for 3 blots, illustrated by red, green and blue)

We note that across all experiments and variables, we found no correlation between 1p-evoked [DA]_o_ levels and the subsequent effect of isradipine (**Figure 1M**, simple linear regression, slope does not significantly differ from zero F_1,28_=0.049 *P* = 0.83) confirming that effect size was not a function of different initial release probability in different conditions.

### Sex differences and calb1 interact to govern LTCC function

VTA and SNc DA neurons differ in a number of key aspects that might contribute to their differing sensitivities to parkinsonian degeneration. One notable difference is the greater expression of calbindin-D28K (Calb1) in VTA neurons, which has been suspected to be protective against PD, although evidence is mixed (Airaksinen et al., 1997; Yuan et al., 2013). We used cre-lox P recombination to selectively knockdown Calb1 in DA neurons in a cross of DAT-Cre mice with floxed Calb1 mice as previously

(Brimblecombe et al., 2019) which does not affect evoked [DA]_o_ or DA uptake in DLS, but it has profound impact in NAc, consistent with the regional endogenous expression profile of Calb1. We found here that in control mice (DAT-Cre hets, referred to as CalbWT), isradipine decreased 1p-evoked [DA]_o_ in males in DLS but not NAc, and did not modify evoked [DA]_o_ in females in DLS (**Figure 2A**) as per wild-type mice (see Fig 1). By contrast, in CalbKD mice, isradipine decreased 1p-evoked [DA]_o_ in male DLS and NAc, and in female DLS (**Figure 2B**). Overall, Calb1 knockdown significantly increased the effect of isradipine on DA release (**Figure 2C**, 2-way ANOVA effect of genotype F_1,14_=31.17 *P<0*.*001*) and enabled LTCCs to support DA release in both male NAc and female DLS. We tested whether there was a gross upregulation of Calb1 levels in the striatum of wild-type female mice relative to males, leading to the occlusion of LTCC function seen in WT female mice. However western blot analysis of protein levels in whole striatal tissue did not detect a difference in Calb1 expression in striatum of females vs males (**Fig 2D,E**, unpaired t-test T_4_=0.18 *P*=0.86).

### DAT function promotes LTCC support of DA release

Translocation of DA across the membrane by DAT involves the co-transport of 2 Na^+^ and 1 Cl^-^, an electrogenic process that can depolarise DA neuron membranes, independently of DA translocation (Ingram et al., 2002) and in cellular expression systems can activate LTCCs (Cameron et al., 2015). We tested whether DATs operate in DA axons to support LTCC function. We inhibited DATs using cocaine (5 μM) or nomifensine (10 μM), monoamine uptake inhibitors which are also known to prevent the electrogenic effects of DAT (Sonders et al., 1997), and tested the impact of isradipine on 1p-evoked [DA]_o_ in DLS in male wild-type mice. We found that in the presence of cocaine or nomifensine, the effect of LTCC inhibitor isradipine in the DLS was no longer apparent (**Figure 3A-C**, Two-way ANOVA: isradipine x pre-condition interaction F_32,192_=3.57, P<0.001; comparison of peaks 1-sample t-test: control t_3_=6.48, P=0.0074, cocaine t_4_=0.37 *P* = 0.73, nomifensine t_4_=1.39 *P*=0.23). The loss of isradipine effect in the presence of cocaine was not due to any potential effects of cocaine as an inhibitor of voltage-gated Na^+^-channels (Na_v_) because application of the Na_v_ blocker and analogue lidocaine did not similarly prevent the effect of isradipine. Rather, in the presence of lidocaine, isradipine decreased 1p-evoked [DA]_o_ as seen in standard conditions (**Figure 3C**, 1-way ANOVA F_2,9_=11.5 *P =* 0.03, Dunnett’s post-test lidocaine vs control *P*= 0.83, cocaine vs control *P* = 0.003, nomifensine vs control *P*= 0.014). These data show that inhibition of DATs occludes LTCCs from contributing to DA release and suggest that DATs support LTCC function in DLS.

**Figure 3.**
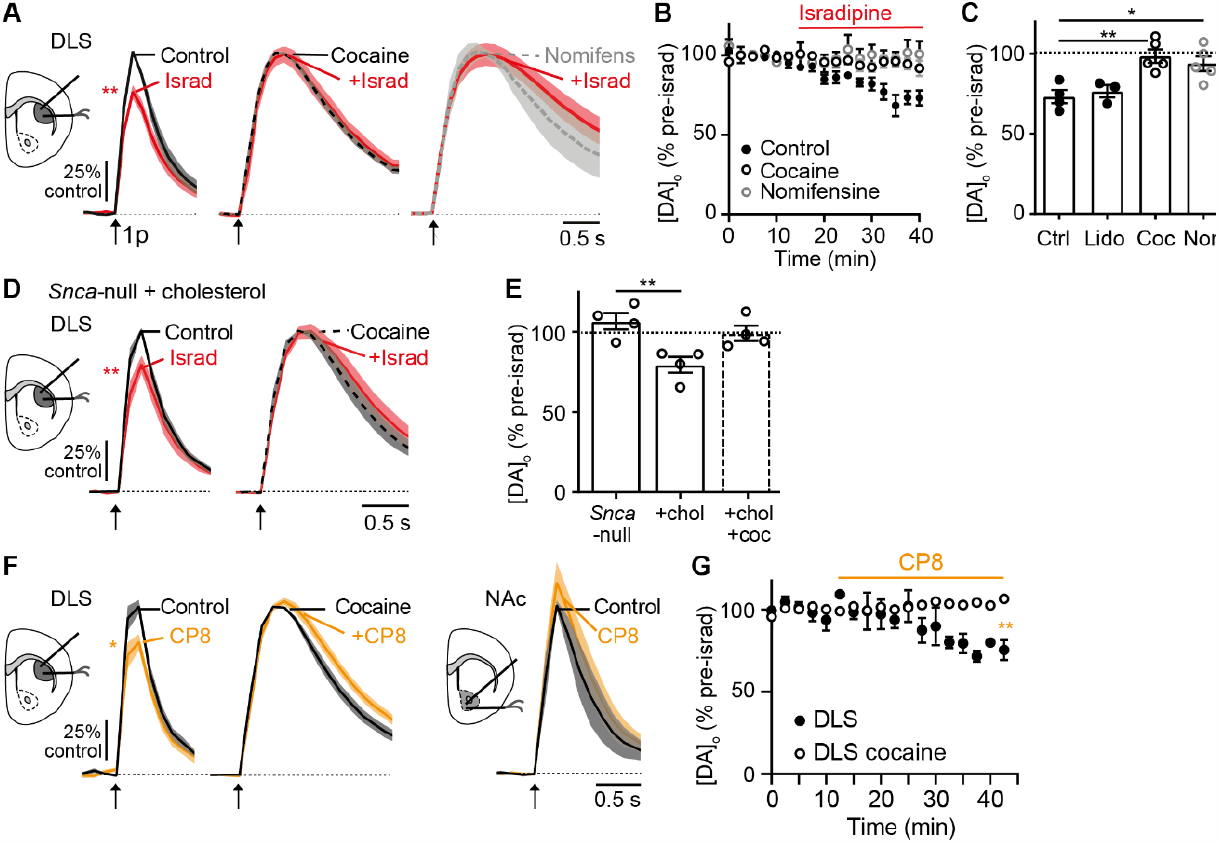
LTCC support of DA release is facilitated by DATs. (**A**) Mean [DA]_o_ ± SEM vs time evoked by single pulses (1p, *arrow*) in male mice in DLS in control versus isradipine (5 μM, *red*) alone (*left*), or in the presence of cocaine (*centre*), or nomifensine (*right*). (**B**) Peak [DA]_o_ ± SEM (normalised to pre-isradipine level) vs time in control (*black*), cocaine (unfilled) or nomifensine (grey unfilled). (**C**) Summary of peak [DA]_o_ remaining after isradipine (% of pre-isradipine control) in DLS of male mice in control (Ctrl) N= 3, lidocaine (*Lido*) N= 3, cocaine (*Coc*) N= 5 or nomifensine (*Nom*) N= 5. (**D**) Mean [DA]_o_ ± SEM vs time evoked by 1p (*arrow*) in DLS in control versus isradipine in *SNCA*-null tissue incubated in cholesterol (*left*) versus cocaine (*right*). (**E**) Summary of peak [DA]_o_ remaining after isradipine (% of pre-isradipine control) in DLS of male *Snca*-null mice in control N= 4, cholesterol incubation N= 4, or cholesterol incubation + cocaine. N= 4 (**F**) Mean [DA]_o_ ± SEM vs time evoked by 1p (*arrow*) in control and CP8 (10 μM, *yellow*) in DLS in control conditions (*left*), in DLS in the prior presence of cocaine (*centre*), or in NAc in control conditions (*right*). (**G**) Peak [DA]_o_ ± SEM (normalised to pre-CP8 level) vs time in DLS in control (*black*) versus cocaine (unfilled), N=4. \**P<*0.05 ** *P*<0.01

We tested whether a potentiation of DAT function could facilitate LTCC support of DA release where it was not previously observable. We have previously shown that DAT function in *Snca*-null mice can be augmented by cholesterol (Threlfell et al., 2021) and exploited this action to see whether in *Snca*-null male mice an effect of isradipine could be brought about after cholesterol incubation and its allied increase in DAT function. Indeed, after incubation in cholesterol, isradipine significantly decreased 1p-evoked [DA]_o_ (**Figure 3D**, control: t_3_=4.2 *P=*0.025). To confirm that this outcome depended on DAT function, we repeated in the presence of cocaine to inhibit DAT. In the presence of cocaine, cholesterol incubation no longer supported an effect of isradipine on DA release (**Figure 3D**, cocaine: t_3_=0.17 *P*=0.87, **Figure 3E**, one-way ANOVA F_3,12_=6.29 *P*= 0.0083, Dunnett’s post-test *Snca*-null control vs cholesterol *P*=0.0034, vs chol+cocaine *P*=0.54), suggesting that a DAT-dependent increase in LTCC function underlay this impact of cholesterol.

We excluded an alternative hypothesis that DAT function promotes the effect of isradipine not by increasing LTCC function but by changing the ability of isradipine to bind to LTCCs. It is known that at hyperpolarised potentials, isradipine does not effectively bind to LTCC (Helton et al., 2005; Rey et al., 2020). Since inhibitors of DATs will reduce DAT-associated depolarising electrogenic currents and maintain DA axons in a more hyperpolarised state, they might prevent isradipine from binding to LTCC. We therefore tested whether the actions of an alternative LTCC antagonist to isradipine, CP8, which binds LTCCs in a voltage-independent manner (Kang et al., 2013; Rey et al., 2020), were also prevented in the presence of cocaine. We found firstly that CP8 decreased 1p-evoked [DA]_o_ in DLS of male mice, but not in male NAc (**Figure 3F,G**, t-test comparison on peak DLS: t_2_=5.19 *P*=0.035, DLS+cocaine t_3_=1.39 *P*=0.26, NAc: t_2_=1.54 *P*=0.26; Two-way ANOVA post-test time point 10 vs 40 min DLS: *P* = 0.0052. NAc (not shown): *P* = 0.78, DLS+cocaine *P =* 0.99), paralleling the effects of isradipine. Further in parallel with the effects of isradipine, CP8 did not reduce 1p-evoked [DA]_o_ in DLS in the presence of cocaine (**Figure 3G**, 2-way ANOVA post-test time point 10 vs 40 min DLS: *P* = 0.0052. NAc: *P* = 0.78, DLS+cocaine *P =* 0.99). These data indicate that DATs indeed support LTCC function in contributing to DA release and do not simply support the voltage-sensitive binding and action of isradipine.

### D2Rs limit LTCC function in NAc

We tested whether D2R activity modifies how LTCCs support DA release by assessing the impact of isradipine on 1p-evoked [DA]_o_ in the presence of D2R inhibition. In DLS, isradipine decreased DA release similarly by ∼25% in the presence and absence of D2R inhibitor L-741,626 (**Figure 4A,B**, comparison of peaks 1 sample t-test control t_3_=6.48 *P*=0.0074; L-741 t_4_=4.87 *P*=0.0082). However in NAc, D2R inhibition unsilenced a role for LTCCs in supporting DA release: it enabled an inhibition of [DA]_o_ by subsequent application of isradipine (**Figure 4C,D**, comparison of peaks 1 sample t-test control: t_3_=1.78, *P*=0.19; L-741: t_3_=27.56 *P*=0.0001), indicating that D2R activity is limiting LTCC function in NAc but not DLS (**Figure 4E**, 2-way ANOVA region X L-741: F_1,14_=8.13, *P=0*.*013*, main effect of region: F_1,14_=16.8, *P=*0.001, main effect of L-741 F_1,14_=14.5 *P*=0.002).

**Figure 4.**
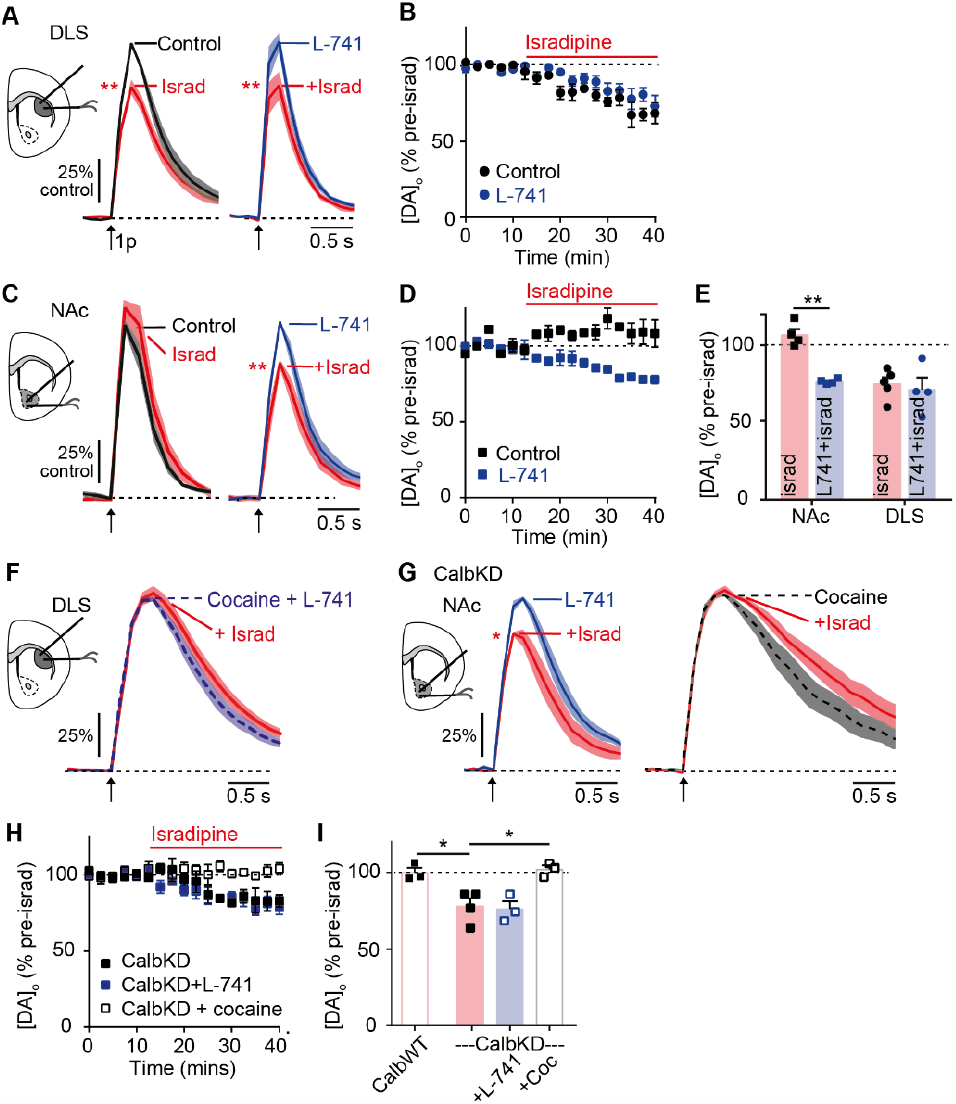
LTCC support of DA release is silenced by D2R. (**A,C**) Mean [DA]_o_ ± SEM vs time evoked by single pulses (1p, *arrow*) in wild-type male mice in DLS (A) or NAc (C) in control versus isradipine (5 μM, *red*) alone (*left*) or in the presence of D2R-antagonist L-741,626 (2 μM, *right*). (**B,D**) Peak [DA]_o_ ± SEM (normalised to pre-isradipine level) vs time in control (*black*) or L-741,626 (*blue*) in DLS (B) or NAc (D). (**E**) Summary of peak [DA]_o_ remaining after isradipine (% of pre-isradipine control), In NAc and CPu without (red) or with L-741 (blue) (N=4) (**F**) Mean [DA]_o_ ± SEM vs time evoked by 1p (*arrow*) in DLS in the presence of cocaine + L-741,626 (*blue dashed*) alone, or with isradipine (*red*). (**G**) Mean [DA]_o_ ± SEM vs time evoked by 1p (*arrow*) in NAc from CalbKD mice with and without isradipine in the prior presence of either L-741,626 (*left*), or cocaine (*right*). (**H**) Peak [DA]_o_ ± SEM (% of pre-isradipine level) vs time during isradipine application in NAc from CalbKD mice in control conditions (*black*), or the presence of L-741,626 (*blue*), or cocaine (*unfilled*). (**I**) Summary of peak [DA]_o_ remaining after isradipine (% of pre-isradipine control) in NAc in CalbWT (N = 3)or CalbKD mice, N=4 with L-741 (N= 3) or cocaine (N= 3). * *P*<0.05 \*\**P*<0.01

**Figure 5:**
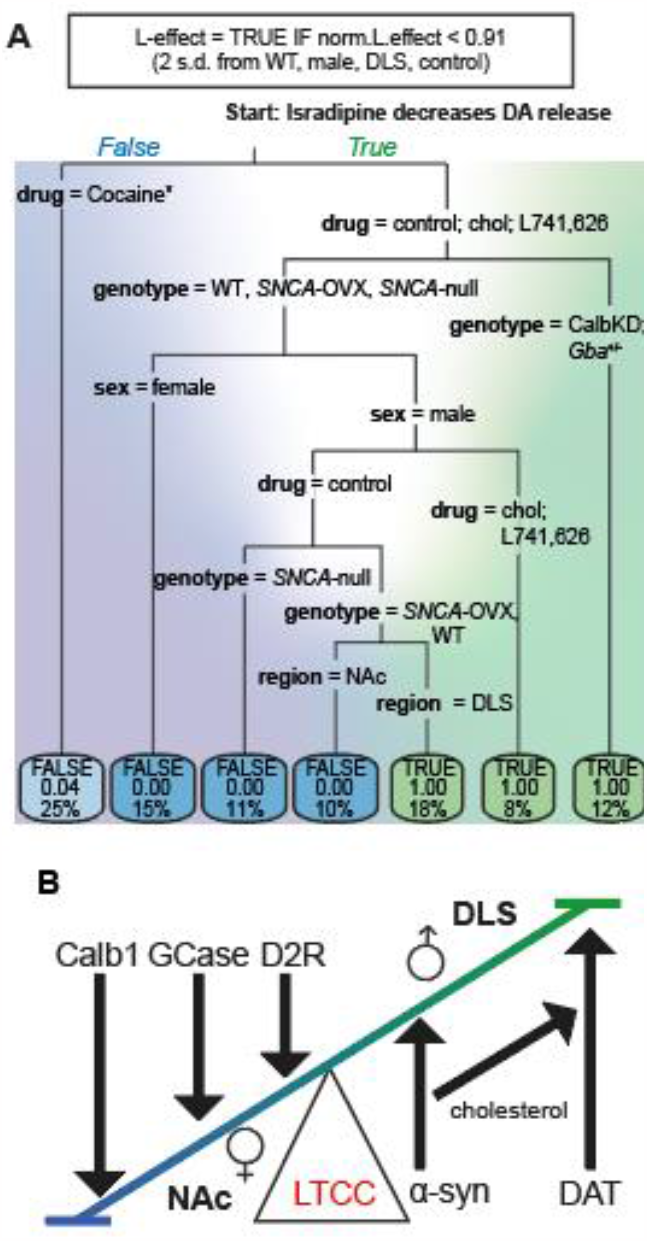
Segregation of factors to facilitators or attenuators of LTCC support of DA release. (**A**) Decision tree segregating experimental conditions of sex, genotype and drug condition using the Gini index. Blue to green background shading shows predictive power of a FALSE (blue) or TRUE (green) outcome respectively. White area shows high uncertainty. Labels on branches indicate recording condition. Left branches for “isradipine decreases DA release” are FALSE. Labels on final boxes (leaves) indicate outcome: TRUE/FALSE (*top*), probability of outcome being correct i.e. 0 or 1 indicate all data from that branch has been correctly sorted (*middle*) and % of datapoints contained within each leaf (*bottom*). *Cocaine is the presence of cholesterol or L-741,626. (**B**) Cartoon illustrating which endogenous factors limit (downward arrows) or promote (upward arrows) LTCC support of DA release.

Given this apparent role for D2Rs in limiting LTCC support of DA release in NAc, we tested a hypothesis that the mechanism through which cocaine limits LTCC function in DLS might be elevation of DA levels leading to a enhancement of D2R-activation. We inhibited D2Rs with L-741,626, to see if this allowed isradipine to decrease evoked [DA]_o_ in the presence of cocaine. However, isradipine did not decrease DA release in DLS in the presence of cocaine and L-741,626, indicating that the control of LTCC support of DA release by DATs in DLS is independent of D2R regulation (**Figure 4F**, t-test t_7_=1.65 P=0.14).

We tested finally whether D2Rs or DATs interact with the effects of Calb1 in NAc on LTCC support of DA release. In NAc of CalbKD mice where LTCCs support DA release (see Fig 2), D2R-inhibition did not augment the impact of isradipine on DA release compared to without D2R inhibition (**Figure 4G-I**, Dunnets post-test t_9_=0.3 *P*=0.94) indicating either that the impact of CalbKD and D2R inhibition of LTCC role has reached a ceiling, or that the mechanism through which D2Rs are governing LTCC impact involves Calb1. The inhibition of DATs in NAc of CalbKD mice with cocaine occluded the effect of isradipine on DA release (**Figure 4G-I**, t_2_=0.73 *P=0*.53). These data indicate that LTCC support of DA release in NAc in CalbKD mice can be facilitated by DATs (**Figure 4I**, 1-way ANOVA Sidak post-test CalbKD vs cocaine t_9_=3.9 *P=0*.*007* CalbKD vs L471 t_9_=0.3 *p=0*.*94*). These features of DAT support but no D2R control of LTCC function in NAc in CalbKD mice mirror the regulation of LTCC function in DLS of wild-type mice. Together, they also reveal that while Calb1 and D2Rs each limit LTCC function in wild-type NAc they do not do so in DLS (in the presence of normal DAT function).

Here, we have shown that the apparent function of LTCCs in supporting DA release varies with a large range of biological conditions, some of which show apparent co-dependencies. To better understand how these factors, of genotype, sex, region and molecular regulators interact to govern whether LTCCs support DA release, and by extension set the conditions when isradipine will decrease DA release, we employed a classification tree algorithm with Gini impurity, for each experimental replicate, to understand the relative influence of each factor (**Figure 5A**). The classification tree illustrated that DAT activity (*cocaine data*) is a key means of supporting L-function; the presence of cocaine is a robust limiter of isradipine’s effect. Conversely, *Calb1* and *Gba* strongly limit LTCC function; knockdown of either promoted isradipine action. In contrast, in wild-type, *SNCA-*OVX and *Snca*-null mice, LTCC function co-varied with sex, striatal region, D2-receptor state or cholesterol level (**Figure 5A**). The branching of the tree into multiple pure leaves (each end node had probability of 0 or 1, no outliers) facilitated a straightforward classification of factors into either facilitators or inhibitors of LTCC function (**Figure 5B**).

## Discussion

We reveal here that LTCC support of DA release from DA axons is regulated by multiple, endogenous factors in a regionally and sexually divergent manner. In DLS, LTCCs operate more evidently in males, and their support of DA release is facilitated by α-synuclein and DATs and permitted by the absence of Calb1. In NAc, LTCC support of DA release is prevented by Calb1 and D2Rs. Intriguingly, regulators that promote LTCC function are associated with increased risk of degeneration in PD, while some of those that limit function are associated with reduced neuronal vulnerability, suggesting that intersecting mechanisms regulating DA handing and LTCC function might together contribute to the metabolic load of vulnerable neurons.

### Sex differences interact with multiple factors to govern LTCC function

LTCCs have been suggested to confer vulnerability to degeneration of SNc DA neurons in PD (Surmeier, 2018), a disease with a higher incidence in men than women (Wooten et al., 2004). Yet, the relative roles of LTCCs in DA neurons in males versus females has been unknown. We reveal there is a greater role for LTCCs in supporting DA release in male DLS than in female, and this raises the speculation that differential LTCC function might contribute to the greater incidence of PD in men than women. Future studies should assess function in both sexes if we are to better understand how to exploit Ca^2+^ biology for a neuroprotective strategy.

Alpha-synuclein is well known to be a contributing factor in PD, however its precise physiological and pathological functions remain incompletely characterised (Alegre-Abarrategui et al., 2019). Here we identify in male mice that axonal LTCC support of DA release is dependent on α-synuclein. It should be noted that some commercially available wild-type mouse strains including the Harlan C57Bl6/6S are null for α-synuclein due to a spontaneous deletion (Specht and Schoepfer, 2001), which likely limits LTCC function in that strain also. The mechanism is yet to be determined, but we note a cascade of potentially interacting factors: we previously found that α-synuclein impacts on brain cholesterols and other lipids that shape DAT activity (Threlfell et al., 2021), and found here that cholesterol reinstates an LTCC function lost after α-synuclein deletion in a DAT-dependent manner. Mutations in the *GBA* gene are the most common genetic factors in PD (Smith and Schapira, 2022) and appear especially important in female patients with PD (Li et al., 2021). Here, we show that *GBA* knockdown led to isradipine decreasing DA release in the DLS of *Gba*^*+/-*^ female mice, indicating that intact GCase activity normally limits how LTCCs support DA release in female mice.

The fast Ca^2+^ buffer calbindin-D28k (Calb1) has been considered previously as potentially neuroprotective, but evidence for a neuroprotective role has not mixed. It remains possible that the toxin models typically used are too aggressive to reveal neuroprotection (Airaksinen et al., 1997; Yuan et al., 2013). In any event, we found that Calb1 was important for both regional and sex differences in LTCC regulation of DA release. Under our standard experimental conditions, LTCCs do not contribute to DA release in male NAc (Brimblecombe et al., 2015), and we show here that Calb1 plays a key limiting role. We found also that Calb1 prevents LTCCs from supporting DA release in DLS of females, despite finding previously that Calb1 does not limit the levels of DA released (Brimblecombe et al., 2019). The mechanisms by which Calb1 limits how LTCCs support DA release have not been fully elucidated and are likely to depend on a number of factors, due to the global signalling nature of Ca^2+^ and the ability of Calb1 to act as a sink and a source for Ca^2+^ (Schwaller et al., 2002; Schwaller, 2020). Potential mechanisms include: Calb1 buffering Ca^2+^ after entry via LTCCs; Calb1 acting as a Ca^2+^ shuttle to facilitate Ca^2+^-dependent inactivation (CDI) of LTCC (Lee et al., 2006); or Calb1 reducing activation of LTCC via inhibition of DAT function (see below). The mechanisms through which Calb1 and LTCC interact in a sex-specific manner are also likely to be complex. Sexual dimorphism in LTCC function has been documented in cardiac tissue (Curl et al., 2008; Prabhavathi et al., 2014; Machuki et al., 2019) whilst in the brain, Calb1 function can be regulated by sex-linked genes and hormones at transcriptional and protein level (Abel and Rissman, 2012). The mechanisms operating in DA neurons are not yet known but could have important clinical implications for sex-specific incidence of disease and responsiveness to medication.

### DAT promotes and D2R limits LTCC support of DA release

We found that LTCC function is supported by DATs, in males in DLS in wild-types and in NAc after calb1 knockdown. Correspondingly, electrogenic monoamine transporters including the DAT are documented to promote Ca^2+^ entry via LTCCs in expression systems (Cameron et al., 2015). We speculate that a similar electrogenic mechanism is responsible in DA axons. It is intriguing to note that in NAc, DA uptake rates are greater after calb1 knockdown than in controls (Brimblecombe et al., 2019), supporting a hypothesis that LTCC support of DA release in NAc after Calb1 knockdown might result from the associated impact of elevated DAT on depolarisation, in addition to a more simple explanation that there is reduced Ca^2+^ buffering. The greater LTCC function in DLS than NAc (Brimblecombe et al., 2015) parallels the higher DAT levels in source DA neurons in SNc than VTA (Poulin et al., 2014). However, DAT function alone is not the sole predictor of LTCC function, as LTCC support of DA release in DLS was not different between wild-type and DAT-Cre^+/-^ mice (CalbWT), despite DAT-Cre^+/-^ mice having lower DAT levels and function (Bäckman et al., 2006; Brimblecombe et al., 2019). Alternative explanations for DAT regulation of LTCC function could yet include that LTCCs might regulate DA release in a DAT-dependent manner, so when DAT is inhibited, LTCC dependence is occluded. Our findings together with other literature add to the growing body of evidence that DATs serve more functions than simply clearing extracellular DA, as they participate in the underlying control of DA release (Venton et al., 2006; Kile et al., 2010; Siciliano et al., 2018; Brimblecombe et al., 2019; Condon et al., 2019).

We also found that LTCC support of DA release is inhibited by D2Rs, in male NAc. LTCCs have vice versa been shown to regulate D2Rs (Zhang and Sulzer, 2012; Dragicevic et al., 2014; Duda et al., 2016), and therefore together these data show that there is reciprocal regulation between LTCCs and D2Rs. D2Rs have been shown previously to inhibit LTCC function in striatal GABAergic projection neurons, and cultured midbrain neurons (Hernández-López et al., 2000; Yasumoto et al., 2004; Olson et al., 2005) but this effect is somewhat surprising for DA release because D2Rs do not appear to be activated sufficiently in this preparation to be limiting DA release for these stimuli (Kennedy et al., 1992; Cragg, 2003; Condon et al., 2019) suggesting that there is limited tonic activity at D2Rs. An explanation for this disparity could be that for 1p-evoked DA release, there is no net effect of inhibiting D2Rs on [DA]_o_ due to the wide range of ion channels and intracellular cascades affected by D2Rs, including voltage-gated K^+^ channel-dependent mechanisms (Fulton et al., 2011), N- and PQ-type Ca^2+^ channels (Cardozo and Bean, 1995) and LTCC-dependent mechanisms (Hernández-López et al., 2000) (reviewed in (Ford, 2014)). These multiple outcomes of D2R activity might also provide an explanation for why D2Rs were not detected to limit LTCC function in DLS. From these collected observations, it is evident that roles of LTCCs in DA release, and the effects of isradipine, are plastic; they are variable between DA neuron types, they vary with the activity of a range of biological regulatory mechanisms, and experimental conditions, and whether the subjects are males or females.

The mechanisms by which LTCCs are supporting DA release, and differently with diverse factors, are not yet resolved. Incoming Ca^2+^ via LTCCs could directly contribute to the Ca^2+^ required for exocytosis or could contribute to other processes that modify DA release probability. Indeed, LTCCs lack the synprint site possessed by several other VGCCs which tethers VGCCs to the exocytotic machinery (Mochida et al., 2003) suggesting LTCCs might not be located in sufficient proximity to catalyse exocytosis directly. Alternatively, LTCCs might support DA release by aiding depolarisation to support AP generation and propagation through the highly branched axonal arbour. They might also couple to Ca^2+^ dependent signalling cascades including those regulated by GPCRs that regulate DA release (Fulton et al., 2011). There is also the possibility that Ca^2+^ entry via LTCCs and downstream activation of RyRs drives mitochondrial generation of ATP (Zampese et al., 2022) that in turn might support axon activity or depolarisation. The factors reported here that change how LTCCs support DA release might change axonal demand for ATP thus driving differing dependencies on Ca^2+^ entry via LTCCs to signal to mitochondria to support the required production of ATP. In turn, since this process contributes a metabolic stress (Pacelli et al., 2015), LTCC function might drive an axonal metabolic stress and also be a reporter for axons with higher metabolic demand. These hypotheses are not mutually exclusive, but are yet to be tested.

In summary, we show that the factors that govern the variable role of axonal LTCCs in supporting DA release in DLS versus NAc include α-synuclein, DATs, calb1 and D2Rs. These factors can vary between striatal regions and support LTCC function in DLS whilst preventing LTCC function in NAc, thereby accounting for the disparity between LTCC expression and function in different DA neurons. It is further noteworthy that factors that promote LTCC support of DA release such as male sex, α-synuclein, DAT and the absence of Calb1 correspond to risk factors for parkinsonian degeneration, understanding how these factors interact may allow the stratification of patient cohorts to identify those who would most benefit from targeting LTCC as a neuroprotective strategy with isradipine (Parkinson Study Group STEADY-PD III Investigators, 2020). By facilitating LTCC function these factors might drive metabolic damage to vulnerable neurons, corroborating the hypothesis that LTCCs might then offer a target for a neuroprotective therapy. Alternatively, LTCC support of DA release might be only a readout of vulnerability, a “canary in the coalmine”, when in turn, manipulation of LTCC function would not protect vulnerable neurons. The utility of targeting LTCC as a neuroprotective strategy against Parkinson’s is likely to depend on there being a net benefit from three competing aspects: firstly, limiting LTCC activity associated with pacemaker function is likely to be beneficial, but only if it outweighs the second factor of reduced dopamine release. Thirdly, it remains to be seen if limiting LTCC associated ATP production is net protective by limiting the production of damaging reactive-oxidative species or damaging, due to it removing a necessary source of ATP. A fuller understanding of the underlying interconnected biology will better inform neuroprotective strategies for PD targeting L-channels and beyond.

## Acknowledgements

The work was funded by grant support from Parkinson’s UK (J-1403; G-1803) and a Clarendon Fund/Christ Church Oxford Studentship to BMR. This research was funded in part by Aligning Science Across Parkinson’s [ASAP-020370] through the Michael J. Fox Foundation for Parkinson’s Research (MJFF).

